# The histone demethylase KDM5C controls female bone mass by promoting energy metabolism in osteoclasts

**DOI:** 10.1101/2023.02.23.529728

**Authors:** Huadie Liu, Lukai Zhai, Ye Liu, Di Lu, Alexandra VanderArk, Tao Yang, Connie M. Krawczyk

## Abstract

Women experience osteoporosis at higher rates than men. Aside from hormones, the mechanisms driving sex-dependent bone mass regulation are not well-understood. Here, we demonstrate that the X-linked H3K4me2/3 demethylase KDM5C regulates sex-specific bone mass. Loss of KDM5C in hematopoietic stem cells or bone marrow monocytes (BMM) increases bone mass in female but not male mice. Mechanistically, loss of KDM5C impairs the bioenergetic metabolism resulting in impaired osteoclastogenesis. Treatment with the KDM5 inhibitor reduces osteoclastogenesis and energy metabolism of both female mice and human monocytes. Our report details a novel sex-dependent mechanism for bone homeostasis, connecting epigenetic regulation to osteoclast metabolism, and positions KDM5C as a target for future treatment of osteoporosis in women.

**One-Sentence Summary:** KDM5C, an X-linked epigenetic regulator, controls female bone homeostasis by promoting energy metabolism in osteoclasts.

## Main Text

Bone mass in adults is controlled by the coordination and balance between osteoclast-mediated bone resorption and osteoblast-mediated bone formation. Osteoclasts (OC) are multinucleated cells of hematopoietic origin formed through the fusion of mononuclear OC precursors from bone marrow monocytes (BMM). Dysregulation of osteoclast-mediated bone resorption has been associated with bone mass related diseases, such as osteoporosis. Women have lower average bone mass than men, conferring a two- to four-fold increase in osteoporosis and age-related fractures (*1*). Systemic pathways—in particular the sex hormones—are prominent in regulating sex-dependent bone mass and have been extensively studied (*2*). However, *ex vivo* cultured female OC progenitors are more potent in osteoclastogenesis than male OC progenitors demonstrating a role for factors intrinsic to OCs regulating sexually dimorphic responses (*3, 4*). Yet the identification of which intrinsic factors that mediate sex-dependent bone homeostasis is still lacking.

Sex chromosomes are the fundamental genetic difference between sexes. Discovering how X- or Y-linked genes contribute to sex-dependent bone mass regulation has the potential to lead to the development of new therapeutics for osteoporosis in women. An increasing number of studies have revealed that osteoclast differentiation and activity is controlled by epigenetic regulation, largely through controlling the accessibility of transcriptional machinery on key osteoclast genes (*5, 6*). However, until now, the epigenetic factors that regulate osteoclast differentiation and function have been found to work similarly in both sexes (*5, 6*). KDM5C (JARID1C/SMCX) is an X-linked <u>lysine</u> H3K4 <u>d</u>e<u>m</u>ethylase that escapes X-inactivation, resulting in higher expression in females than males (*7*). Global loss of KDM5C results in spurious transcription of genes normally silenced during development (*8, 9*). In men and male mice, variants of KDM5C cause X-linked intellectual disability (XLID) with short stature, aggressive behavior, and autism (*10–12*). Despite the short stature observed in these individuals, a role for KDM5C in bone homeostasis has not been reported. Herein, we report that the loss of KDM5C in BMM increases bone mass in female mice, impairs osteoclastogenesis, and reduces bioenergetic metabolism in osteoclast precursors. Thus, KDM5C represents a cell-intrinsic, sex-dependent epigenetic regulator of osteoclastogenesis and bone mass regulation in females, and its inhibition provides a potential therapeutic strategy for preventing osteoporosis in females.

## Results

### Loss of KDM5C in hematopoietic cells results in increased trabecular bone mass in female mice

To investigate the function of KDM5C in the hematopoietic-osteoclast lineage, we generated *Vav-iCre*; *Kdm5c*^fl/fl^ mice, which diminishes KDM5C expression in all hematopoietic cells including BMM that give rise to osteoclasts. No severe global developmental defects were observed in either sex of the KDM5C-conditionally deficient mice. However, upon closer examination, we noticed a marked increase in macroscopic trabecular bone volume in female *Vav-iCre*; *Kdm5c*^fl/fl^ mice (f*Kdm5c*^ΔVav^). When analyzed for bone density and microstructure by MicroCT, bones from f*Kdm5c*^ΔVav^ mice were found to have significantly increased trabecular bone mass at 16 weeks of age (**Fig. 1A**), as indicated by increased trabecular bone volume/ total volume (BV/TV), trabecular number (Tb.N), trabecular thickness (Tb.Th), and decreased trabecular separation (Tb.Sp) (**Fig. 1B**) compared to female control littermates (fCtrl). In contrast, the male *Vav-Cre*; *Kdm5c*^fl/fl^ mice (m*Kdm5c*^ΔVav^) and female KDM5C heterozygous conditional knockout mice (f*Kdm5c*^ΔVav^) have no obvious trabecular bone architecture difference compared to mCtrl and fCtrl, respectively (**Fig. 1A, B**). Parameters of cortical bone did not show significant changes between groups (**Fig. 1C**). These data reveal a sex-dependent role for KDM5C in regulating trabecular bone mass.

**Fig. 1.**
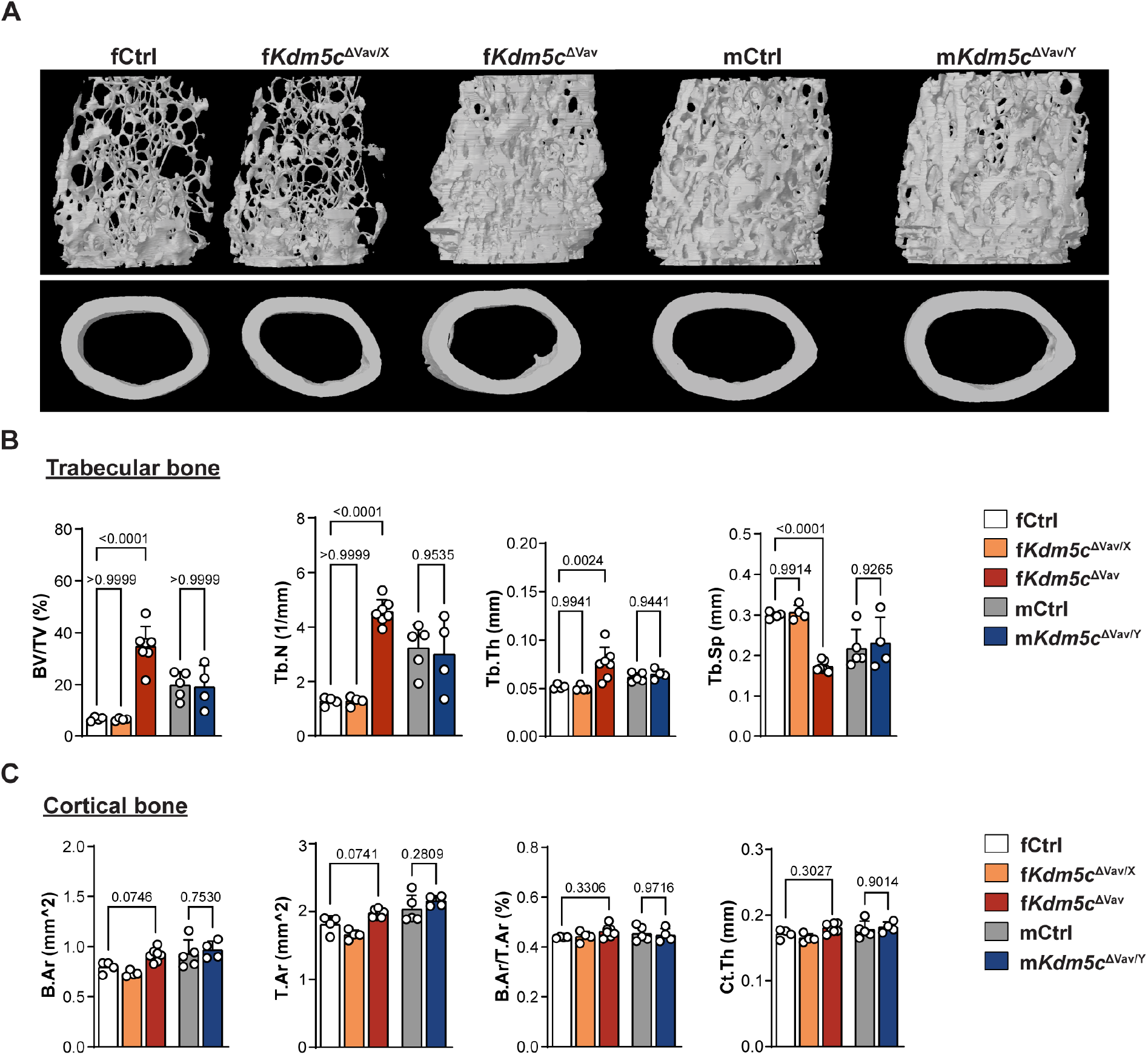
Increased trabecular bone mass in distal femurs of female *Kdm5c*^ΔVav^ but not male *Kdm5c*^ΔVav^ mice. (**A**) Representative micro-CT images and (**B**, **C**) quantitation of femur trabecular bone (upper panel) and cortical bone (lower panel) from 16-week-old mice of indicated genotypes/sex. Data comparisons are conducted using one-way ANOVA analyses. Each data point represents an individual mouse. Data are presented as mean ± s.e.m.

### KDM5C intrinsically regulates osteoclastogenesis

Bone mass in adults is regulated by the coordination and balance between osteoclast-mediated bone resorption and osteoblast-mediated bone formation. We examined bone slices for the presence of OC using tartrate-resistant acid phosphatase (TRAP) staining. We found that the bones from fKdm*5c*^ΔVav^ mice have significantly decreased osteoclast surface/bone surface (Oc.S/BS) and osteoclast number/bone surface (N.Oc/BS) in trabecular bone compared to controls (**Fig. 2A**), indicating that osteoclastogenesis and bone resorption is impaired. Other hematopoietic cell types such as T cells and B cells that participate in bone mass regulation indirectly, by affecting the bone microenvironment (*13*), would also be affected in the *Vav-iCre* mice. To elucidate whether KDM5C regulates bone formation intrinsically in the myeloid lineage, including monocytes and osteoclasts, we isolated BMMs from bone marrow of control and f*Kdm5c*^ΔVav^ mice and analyzed their ability to generate osteoclasts *ex vivo* following RANKL treatment. Osteoclast formation was significantly impaired in f*Kdm5c*^ΔVav^ BMMs, as indicated by the reduced osteoclast number (Oc.N) and osteoclast surface (Oc.S) (**Fig. 2B**), and decreased expression of osteoclastogenic gene transcripts (**Fig. 2C**). Furthermore, we generated *LysM-Cre*; *Kdm5c*^fl/fl^ mice which have KDM5C deleted in myeloid cells. Similar to f*Kdm5c*^ΔVav^ mice, female *LysM-Cre*; *Kdm5c*^fl/fl^ mice (f*Kdm5c*^ΔLysM^) had significantly increased trabecular bone mass and decreased osteoclast activity, as indicated by increased BV/TV and Tb.N; decreased Tb.Sp, Oc.S/BS, and N.Oc/BS (**Fig. 3A, B**). Consistent with the f*Kdm5c*^ΔVac^ mice, we observed no significant differences in cortical bone mass (**Fig. 3A**). Overall, the bone phenotypes in the f*Kdm5c*^ΔLysM^ were less severe compared to the f*Kdm5c*^ΔVav^ mice. We found that BMM isolated from f*Kdm5c*^ΔLysM^ mice show significant reduction in osteoclastogenic potential *ex vivo,* however not to the same extent as observed in the *Kdm5c*^ΔVav^ BMM (**Fig. 3C-D**). This is likely due to reduced KDM5C deletion efficiency in f*Kdm5c*^ΔLysM^ BMM compared to f*Kdm5c*^ΔVav^ BMM (**Fig. S1A-D**). Together, our findings demonstrate that loss of KDM5C intrinsically in myeloid progenitor cells results in reduced osteoclastogenesis and lower bone mass, in female mice only.

**Fig. 2.**
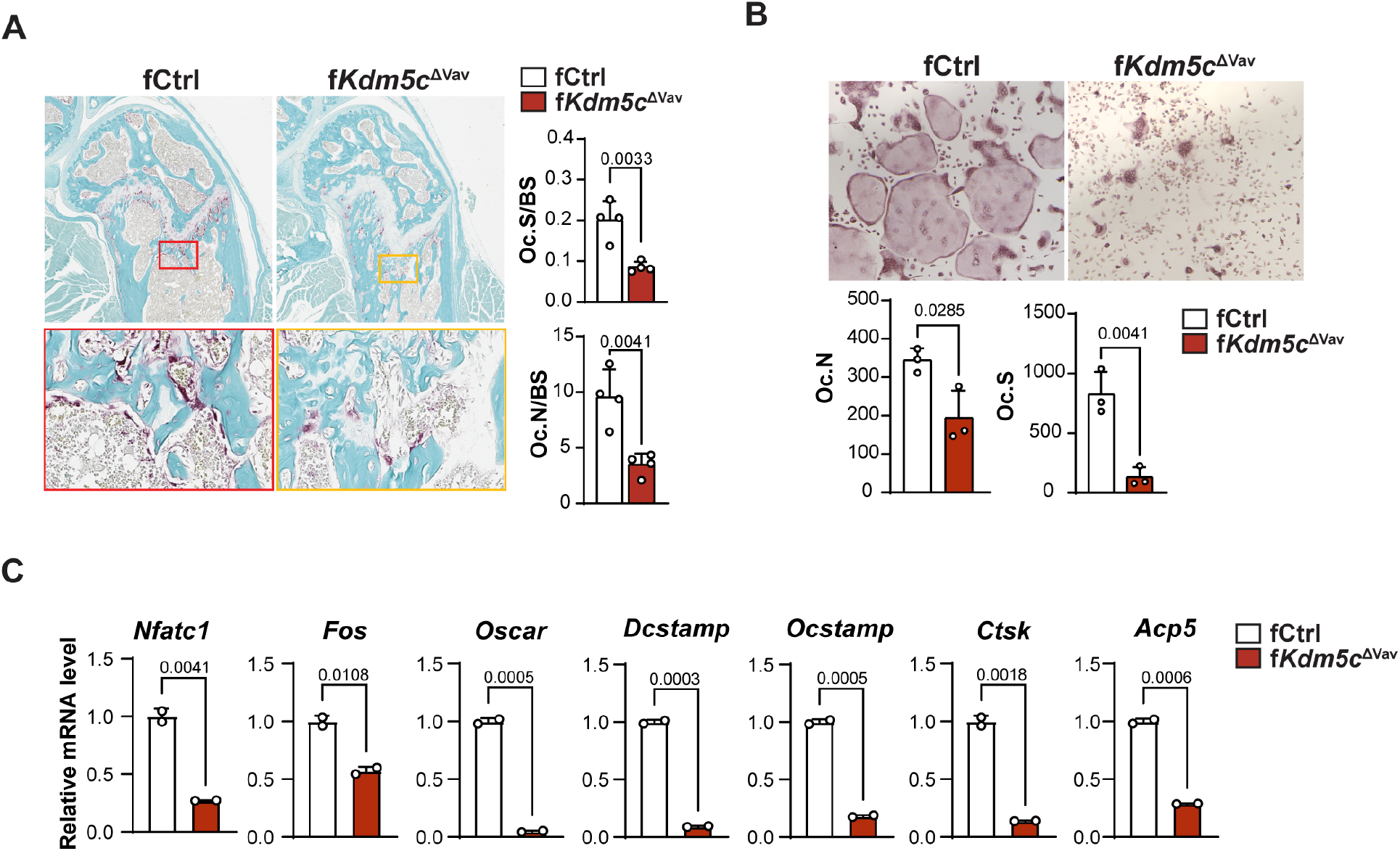
Impaired osteoclastogenesis in female *Kdm5c*^ΔVav^ mice. (**A**)TRAP staining of sectioned femurs from 16-week-old fCtrl and f*Kdm5c*^ΔVav^ mice. OC number/bone surfaces (Oc.N/BS) and OC surfaces/bone surfaces (Oc.S/BS) in trabecular bone area were measured and calculated (*n*=4 per genotype). (**B**) *ex vivo* osteoclastogenesis on BMMs of fCtrl and f*Kdm5c*^ΔVav^ mice. OCs were visualized by TRAP staining 3-5 days after differentiation. Oc.N and Oc.S were measured and calculated (*n*=3 per genotype). (**C**) mRNA levels of osteoclast/osteoclastogenic genes in fCtrl and f*Kdm5c*^ΔVav^ BMM 48hrs after osteoclastogenic induction. mRNA levels were detected by qRT-PCR. Relative mRNA levels are shown (*n*=2 per genotype). All data comparisons are conducted by Student’s *t*-test, two-tailed. Data are presented as mean ± s.e.m.

**Fig. 3.**
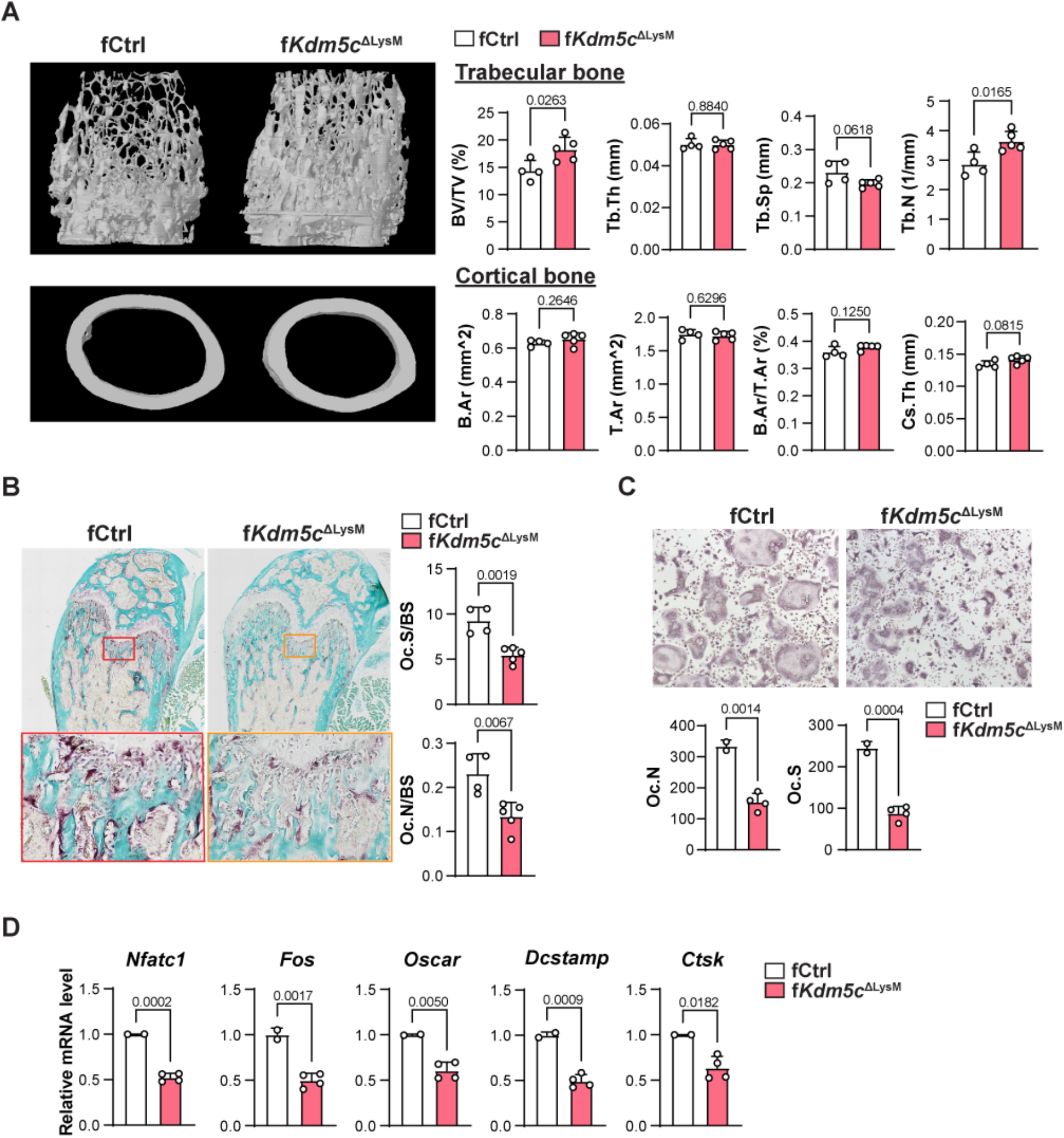
Increased trabecular bone mass and impaired osteoclastogenesis in f*Kdm5c*^ΔLysM^ mice. (**A**) Representative micro-CT images and quantitation of femur trabecular bone and cortical bone from 8-week-old fCtrl and f*Kdm5c*^ΔLysM^ mice. (**B**) Representative TRAP staining images of sectioned femurs from 8-week-old fCtrl and f*Kdm5c*^ΔLysM^ mice, used to calculate OC number/bone surfaces (Oc.N/BS) and OC surfaces/bone surfaces (Oc.S/BS) (*n*=4 for fCtrl; *n*=5 for f*Kdm5c*^ΔLysM^). (**C**) *ex vivo* osteoclastogenesis of BMMs from fCtrl and f*Kdm5c*^ΔLysM^ mice. OCs were visualized by TRAP staining 3-5 days after differentiation. Oc.N and Oc.S were measured and calculated (*n*=3 per genotype). (**D**) mRNA levels of osteoclast/osteoclastogenic genes were detected by qRT-PCR in fCtrl and f*Kdm5c*^ΔVav^ BMM 48hrs after induction. Relative mRNA levels are shown (*n*=2 for fCtrl, *n*=4 for f*Kdm5c*^ΔLysM^). All data comparisons are conducted by Student’s *t*-test, two-tailed. Data are presented as mean ± s.e.m.

### Mitochondrial respiration and energy metabolism is impaired following the loss of KDM5C in BMMs

KDM5C represses transcription via demethylation of H3K4me2/3 at gene promoters, but also promotes gene expression in mouse embryonic stem cells (*9, 14*) and ERα-positive breast cancer cells (*15*) by converting H3K4me2/3 modifications into H3K4me1 or recruiting transcription factors on specific transcriptional enhancers. Therefore, to investigate how KDM5C affects transcriptional programming of BMM, we compared gene expression of fCtrl and f*Kdm5c*^ΔVav^ BMMs at different stages of osteoclastogenesis *ex vivo* (0h, 16h, and 32h). Significant transcriptomic differences were observed between fCtrl and f*Kdm5c*^ΔVav^ at all three stages (**Fig. S2**). Using Gene Ontology analysis, we found genes that were increased in KDM5C-deficient BMM/OCs were related to immune cell activity and inflammation, while genes with decreased expression were enriched in metabolic and mitochondrial respiration-related pathways (**Fig. 4A**). Notably, mitochondrial ATP synthesis and electron transport pathways are repressed the most in KDM5C-deficient cells at early stages of osteoclast formation (**Fig. 4A**, 32h). These data demonstrate that KDM5C positively regulates cell metabolism in BMM/OC.

**Fig. 4.**
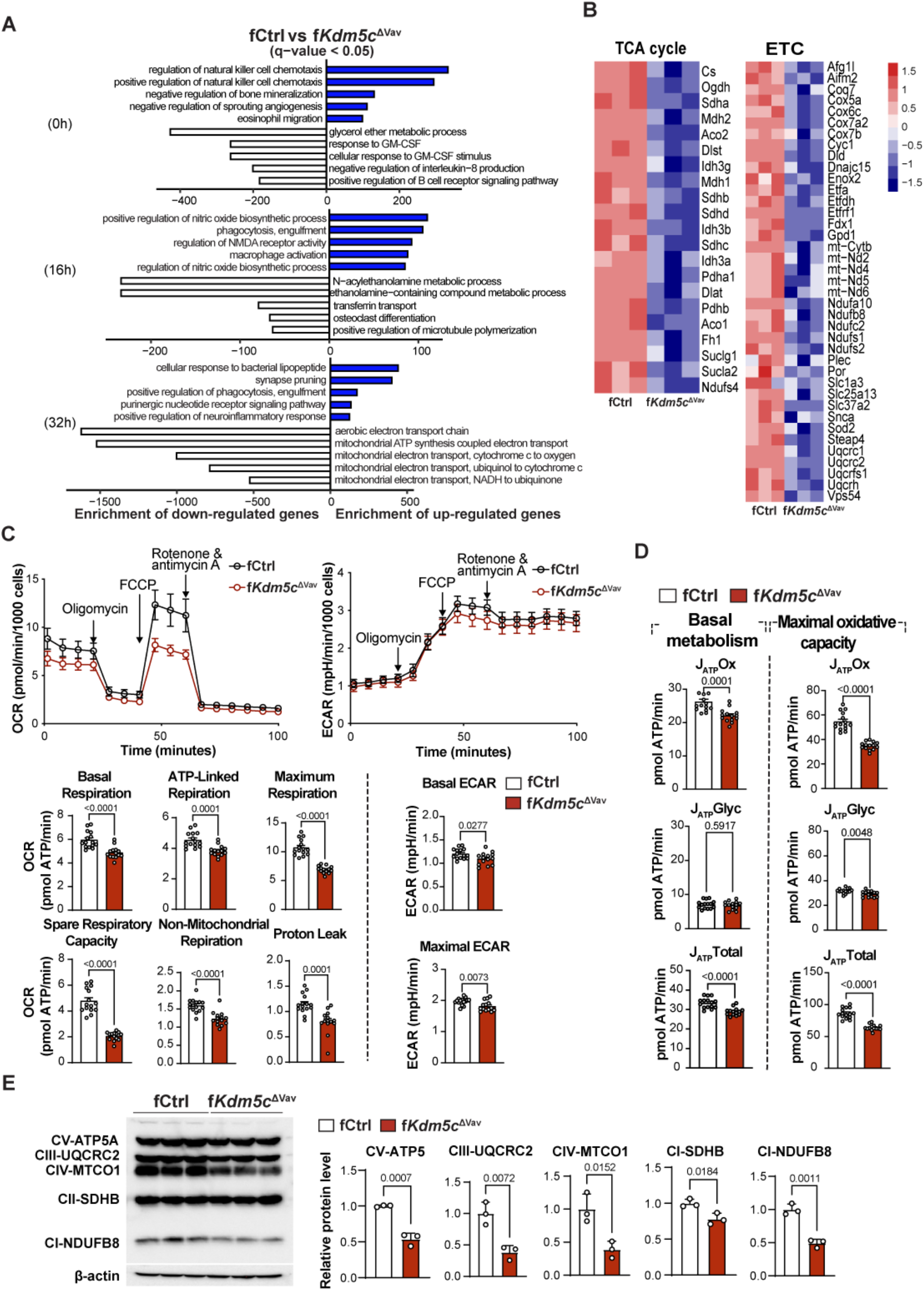
KDM5C-deficient BMMs have decreased bioenergetic metabolism during osteoclastogenesis. (**A**) The top 5 enriched gene ontology biological process (GOBP) terms assigned to up-regulated (blue bars) or down-regulated (white bar) genes in f*Kdm5c*^ΔVav^ vs fCtrl cells during osteoclastogenesis. Gene expression was determined by RNA-seq at three time points during osteoclastogenesis. Genes with FDR<0.05 were chosen for analysis. (**B**) Heatmap of TCA cycle genes and mitochondrial electron chain (ETC) genes expression in f*Kdm5c*^ΔVav^ and control BMMs 32h after osteoclastogenic induction. (**C**) Oxygen consumption rate (OCR, left) and extracellular acidification rate (ECAR, right) in fCtrl and f*Kdm5c*^ΔVav^ BMMs. OCR and ECAR were detected 72 hours after osteoclastogenic induction. Detailed parameters including basal and maximum respiration, ATP-coupled respiration, spare respiratory capacity, non-mitochondrial respiration, and proton leak are shown in the bottom panels. (**D**) Oxidative ATP production rate and maximum glycolytic ATP production in f*Kdm5c*^ΔVav^ and control BMMs. (**E**) Immunoblot of mitochondrial OXPHOS complex proteins (*left)* and densitometry quantifications of control and *Kdm5c*-deficient BMMs stimulated by RANKL for 72 hours. Each lane represents one mouse (*n*=3 per genotype). All data comparisons are conducted by Student’s *t*-test, twotailed. Data are presented as mean ± s.e.m.

Osteoclastogenesis and osteolysis are energy-consuming processes (*16*), and preosteoclast differentiation and osteoclast survival are suppressed when mitochondrial metabolism is impaired (*16–18*). Because TCA cycle and mitochondrial genes were decreased in KDM5C-deficient BMM (**Fig. 4B**), we hypothesized that KDM5C augments osteoclastogenesis by promoting mitochondrial metabolism in BMM. We measured oxygen consumption rate (OCR), and extracellular acidification rate (ECAR) as surrogates of mitochondrial metabolism and glycolysis, respectively (**Fig. 4C**). Notably, basal respiration, ATP-coupled respiration, spare respiratory capacity, and maximal respiratory capacity were all significantly reduced in KDM5C-deficient BMM compared to controls (**Fig. 4C**). We also found that KDM5C-deficient BMM has reduced glycolysis (**Fig. 4C**). These results demonstrate that KDM5C positively regulates bioenergetic metabolism. This was confirmed by the decrease in ATP production from oxidative phosphorylation (OXPHOS) and glycolysis (**Fig. 4D**). Our results also confirm previous results that osteoclasts generate most of their ATP via OXPHOS (*17*) (**Fig. 4D**). We examined the expression of proteins involved in electron transport chain (ETC) assembly and function and found that ATP5, UQCRC2, MTCO1, SDHB, and NDUFB8 were all reduced in the KDM5C-deficient cells relative to controls (**Fig. 4E, Fig. S3A**). Similarly, genes involved in ETC assembly and function including *Atp6vod2, Uqcrc2, Atp1b3, Cox6b1, Sdhb,* and *Sdha* had lower expression in KDM5C-deficent BMM (**Fig. S3B**). Overall, these data demonstrate that KDM5C is necessary for bioenergetic metabolism in BMM/OC, largely through promoting mitochondrial respiration.

### KDM5C-regulated osteoclastogenesis is mediated in part by PGC-1β

Next, we investigated how KDM5C loss alters metabolic programming in the BMM/OC population. PGC-1α and PGC-1β are transcriptional co-activators that promote mitochondrial biogenesis and the expression of mitochondrial metabolism genes (*19*). Previous studies reported that PGC-1β supports osteoclast formation (*20, 21*) by enhancing mitochondrial biogenesis and cytoskeletal rearrangement (*22*). We found that PGC-1β (*Ppargc1b*), but not PGC1-α (*Ppargc1a*), was highly expressed in control BMM and increased during osteoclastogenesis (**Fig. 5A**). We found that PGC-1β expression did not increase to the same extent in KDM5C-deficient BMM/OC (**Fig. 5A** and **Fig. S3C**). To determine if increasing the expression of PGC-1β in KDM5C-deficient BMM could rescue osteoclastogenesis, we overexpressed PGC-1β in control and KDM5C-deficient BMMs using retroviral expression. We found that the osteoclastogenic potential of KDM5C-deficient BMMs could be partially restored by PGC-1β, as indicated by increased Oc.N and Oc.S in an *ex vivo* osteoclastogenesis assay (**Fig. 5B**). PGC-1β-mediated regulation of mitochondrial metabolism has been shown to be dependent on iron uptake by the Transferrin Receptor protein 1, TfR1(*20*), and iron itself is required for synthesis of cofactors essential to the function of enzymes in the electron transport chain. We examined the expression of *Tfrc* (the gene encoding TfR1) in KDM5C-deficient BMM/OC and found decreased expression relative to controls (**Fig. S4**). These data show that PGC-1β mediates KDM5C-regulated osteoclastogenesis, but other factors, including iron uptake via TfR1, and/or the expression of other mitochondrial genes are also likely contributing.

**Fig. 5.**
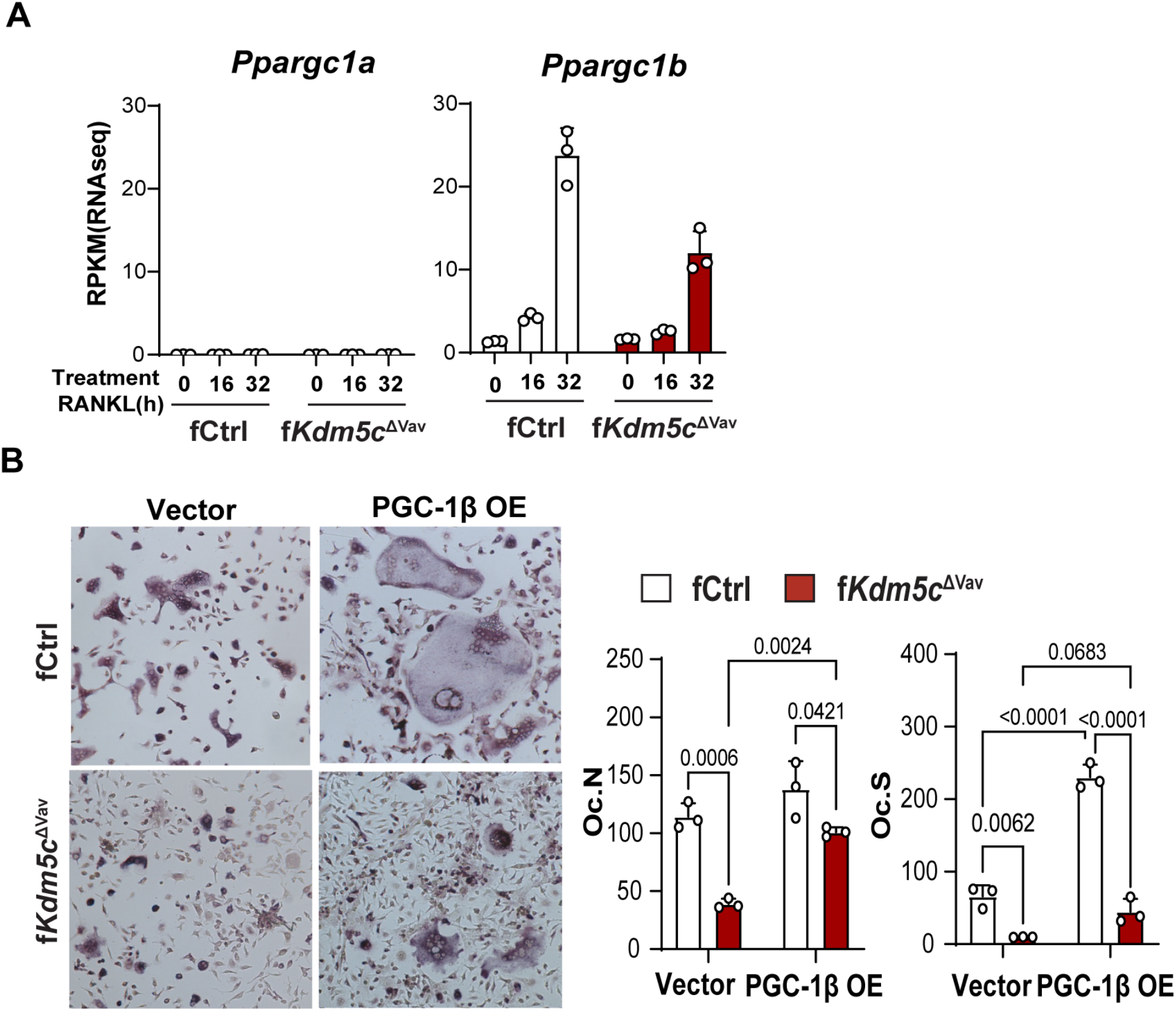
Partial rescue of osteoclastogenesis by PGC1-b. (**A**) *Ppargc1b* and *Ppargc1a* expression (RNA-seq) during *ex vivo* osteoclastogenesis assay at indicated time points. (**B**) *ex vivo* osteoclastogenesis measured by TRAP staining of BMMs from fCtrl and f*Kdm5c*^ΔVav^ mice transduced with control or PGC1-b overexpression vector (two-way ANOVA analyses). Data are presented as mean ± s.e.m.

### KDM5 inhibition dampens osteoclastogenic potential of both mouse and human monocytes

Next, we tested whether we could prevent osteoclastogenesis in fCtrl BMMs by using small molecule inhibitors of KDM5C demethylase activity. While no KDM5C specific inhibitor exists, pan-KDM5 inhibitors (KDM5i) are available and have been used in clinical trials for the treatment of cancer and hepatitis B (*23, 24*). Our RNA-seq data shows that *Kdm5c* is expressed more highly than other KDM5 family members in female BMM (**Fig. 6A**), suggesting that KDM5C may have a greater role than other KDM5 members in regulating osteoclastogenesis. We found that KDM5i dose-dependently suppressed RANKL induced osteoclastogenesis of female BMMs, as indicated by TRAP staining in an *ex vivo* osteoclastogenesis assay, with an IC50 of 5.6 μM (**Fig. 6B**). Consistently, KDM5i treatment significantly downregulated key mitochondria OXPHOS complex proteins (**Fig. 6C)** and mRNAs (*Sdhb, Atp6v0d2, Sdha, Uqcrc2, Uqcrfs1*), as well as mRNA expression of osteoclast marker genes (*Dcstamp, Acp5, Ctsk*) and *Ppargc1b* (**Fig. S5A**). OCR, ECAR, and ATP production were also impaired by KDM5i in a dose dependent manner (**Fig. 6D, E**), demonstrating that the function of KDM5C in regulating mitochondrial metabolism is dependent on its demethylase activity. Importantly, KDM5i increased H3K4me3 level in CD14+ monocytes of human PBMC (**Fig. S5B)**. The effect of KDM5i on osteoclastogenesis and energy metabolism are conserved in female mice and humans, indicated by the reduction in the number and area of multinuclear osteoclasts (**Fig. 6F**), and the impairment in OCR, ECAR, and ATP production (**Fig. 6G** and **Fig. S5C**) in KDM5i treated female peripheral human blood monocytes under osteoclastogenic differentiation. Thus, targeting KDM5C has the potential to reduce osteoclast function therapeutically, potentially mitigating bone loss in females.

**Fig. 6.**
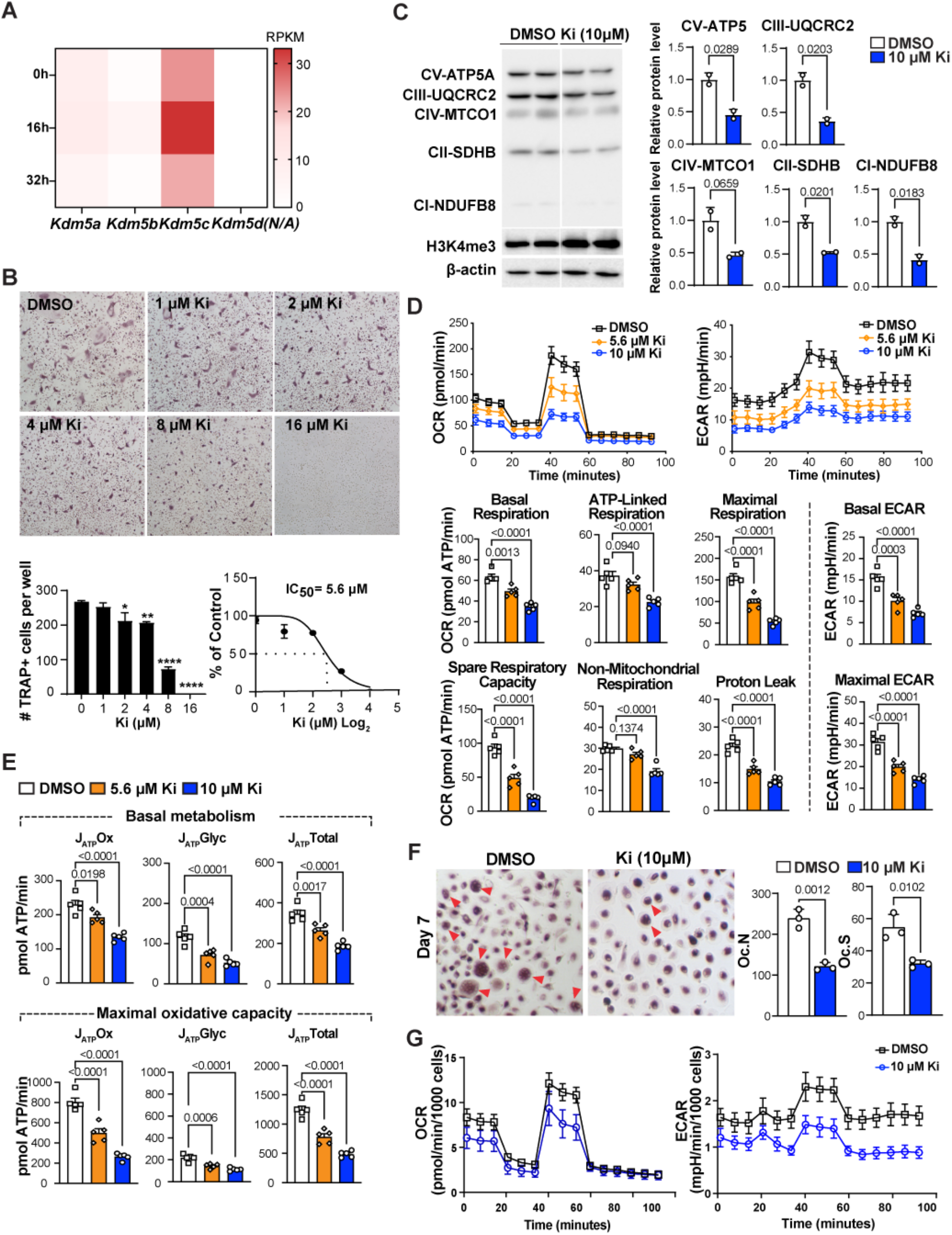
Pharmacological inhibition of KDM5 impairs mitochondrial metabolism and osteoclastogenesis of mice and human monocytes. (**A**) The gene expression of KDM5 family members during osteoclastogenesis (analyzed by RNAseq, as in Fig. 5A). (**B**) Representative TRAP staining images of BMM osteoclastogenesis following treatments with different doses of KDM5 inhibitor (Ki) (Top). The number of TRAP positive cells per well were quantified and used to calculate IC50 of KDM5i in osteoclastogenesis (Bottom). (**C**) Western blots of mitochondrial OXPHOS proteins and H3K4me3 after Ki (10 μM) treatment. (**D**) OCR (left) and ECAR (right) of RANKL-stimulated (72h) BMM treated with DMSO, 5.6 μM or 10 μM Ki. All parameters were calculated as in Fig. 4. (**E**) Oxidative and glycolytic ATP production of BMM following Ki treatment. (**F**) Human PBMC osteoclastogenesis with Ki treatment. Monocytes cultured from human PBMC were treated with RANKL for 7 days before TRAP staining. Multinuclear (>3) osteoclasts are indicated by red arrows. Total numbers (Oc.N) and total area (Oc.S) of osteoclasts were quantified using ImageJ (*n=3* for each group). (**G**) OCR (left) and ECAR (right) of human PBMC-monocytes treated with DMSO and 10 μM Ki 5 days after osteoclastogenic induction. Comparisons in C and F are conducted by Student’s *t*-test, twotailed; in D and E by one-way ANOVA analyses. Data are presented as mean ± s.e.m.

## Discussion

Our work has identified KMD5C, an X-linked chromatin modifying enzyme, as a female-specific regulator of bone mass that inhibits OC differentiation and function. Females, without any KDM5C in BMM/OC displayed increased bone mass comparable to control or KDM5C-deficient males demonstrating that KDM5C uniquely regulates bone mass in females. Mechanistically, we found that KDM5C promotes bioenergetic metabolism required for osteoclastogenesis. Thus, we have identified a new mechanism linking epigenetic and metabolic programming of OCs to sex-specific bone mass regulation. Females are at a higher risk than males for developing osteoporosis, mainly because of the inherent lower peak bone mass before menopause, and a more aggressive bone loss at post-menopause stage. Estrogen inhibits osteoclast formation and function and was used for osteoporosis treatment for women, but also leads to strong side effects, including cancers. Current treatment alternatives to estrogen have been developed and successfully used to treat osteoporosis, however none of these therapeutics target a mechanism specific to females. Our study showed that pharmacological inhibition of KDM5C blocks osteoclastogenesis of both human and mouse monocytes, indicating that KDM5C is a viable therapeutic target for osteoporosis, particularly for females.

Our data show that KDM5C promotes mitochondrial metabolism and related gene expression. While KDM5C predominantly functions as a transcriptional repressor by removing active H3K4me2/3 marks from promoters, it can also stimulate gene expression dependent or independent of demethylase activity(*9, 14, 15*). In our study, use of the KDM5-specific histone lysine demethylase inhibitor reduces mitochondrial metabolism and related gene expression, demonstrating that KDM5C demethylase activity is critical for BMM/OC bioenergetic metabolism. Demethylase-dependent positive regulation of gene expression by KDM5C occurs by trimming H3k4me2/3 for optimal enhancer activity (*9*). Thus, KDM5C may control the mitochondrial transcriptional program in osteoclasts through enhancer regulation. In particular, *Ppargc1b,* which encodes PGC-1β, a master regulator of mitochondrial metabolism and biogenesis, is significantly downregulated. While we found that PGC-1β ectopic expression partially rescued osteoclastogenesis, PGC-1β is not a rate limiting mediator of KDM5C-mitochondrial metabolism on its own. Notably, expression of the iron transporter TfR1 was reduced in the absence of KDM5C; this suggests that the KDM5C-dependent iron uptake mechanism and PGC-1β expression may synergize to promote mitochondrial metabolism and osteoclastogenesis (*20*).

Overall, our findings highlight a female-specific epigenetic mechanism that controls osteoclastogenesis by governing the transcriptional programming of energy metabolism, positioning KDM5C as a novel target for the treatment of osteoporosis in females.

## Acknowledgments

We thank the current and former members of C. M. Krawczyk and T. Yang laboratories and Bart Williams for thoughtful discussions, and Drs. Sonya Craig and Carolyn Anderson for editing. We are indebted to Dr. Anding Shen from Calvin University for providing human PMBC. This work was supported by the excellent staff in the Vivarium and Transgenics, Genomics, Pathology & Biorepository, and Metabolomics and Bioenergetics Cores at Van Andel Institute. This work was supported by the Van Andel Institute Employee Impact Fund for Research awarded to TY and CK (2021), VAI startup funds (C.K., and T.Y.) and the NIH (T.Y.; R01AR079174).

## Author contributions

HL, LZ, TY, and CMK conceived this study and designed experiments. HL, LZ, DL, YL, AVA executed experiments and performed data analysis. HL, LZ, YL, and DL generated figures. HL and LZ wrote the manuscript, and TY and CMK edited the manuscript.

## Data and materials availability

All data are available in the main text or the supplementary materials.

## Supplementary Materials

Materials and Methods

Figs. S1 to S5

References (*25–31*)

## References and Notes

1. O. Lofman, L. Larsson, G. Toss, Bone mineral density in diagnosis of osteoporosis: reference population, definition of peak bone mass, and measured site determine prevalence. J Clin Densitom 3, 177–186 (2000).

2. S. Khosla, D. G. Monroe, Regulation of Bone Metabolism by Sex Steroids. Cold Spring Harb Perspect Med 8, (2018).

3. S. H. Mun et al., Sexual Dimorphism in Differentiating Osteoclast Precursors Demonstrates Enhanced Inflammatory Pathway Activation in Female Cells. J Bone Miner Res 36, 1104–1116 (2021).

4. D. N. Paglia et al., Runx1 Regulates Myeloid Precursor Differentiation Into Osteoclasts Without Affecting Differentiation Into Antigen Presenting or Phagocytic Cells in Both Males and Females. Endocrinology 157, 3058–3069 (2016).

5. K. Astleford, E. Campbell, A. Norton, K. C. Mansky, Epigenetic Regulators Involved in Osteoclast Differentiation. Int J Mol Sci 21, (2020).

6. F. Xu et al., The Roles of Epigenetics Regulation in Bone Metabolism and Osteoporosis. Front Cell Dev Biol 8, 619301 (2020).

7. P. A. Lingenfelter et al., Escape from X inactivation of Smcx is preceded by silencing during mouse development. Nat Genet 18, 212–213 (1998).

8. M. Scandaglia et al., Loss of Kdm5c Causes Spurious Transcription and Prevents the Fine-Tuning of Activity-Regulated Enhancers in Neurons. Cell Rep 21, 47–59 (2017).

9. N. S. Outchkourov et al., Balancing of histone H3K4 methylation states by the Kdm5c/SMCX histone demethylase modulates promoter and enhancer function. Cell Rep 3, 1071–1079 (2013).

10. S. Iwase et al., A Mouse Model of X-linked Intellectual Disability Associated with Impaired Removal of Histone Methylation. Cell Rep 14, 1000–1009 (2016).

11. F. Abidi et al., Novel human pathological mutations. Gene symbol: JARID1C. Disease: mental retardation, X-linked. Hum Genet 125, 345 (2009).

12. F. E. Abidi et al., Mutations in JARID1C are associated with X-linked mental retardation, short stature and hyperreflexia. J Med Genet 45, 787–793 (2008).

13. M. N. Weitzmann, T-cells and B-cells in osteoporosis. Curr Opin Endocrinol Diabetes Obes 21, 461–467 (2014).

14. M. K. Samanta et al., Activation of Xist by an evolutionarily conserved function of KDM5C demethylase. Nat Commun 13, 2602 (2022).

15. H. F. Shen et al., The Dual Function of KDM5C in Both Gene Transcriptional Activation and Repression Promotes Breast Cancer Cell Growth and Tumorigenesis. Adv Sci (Weinh) 8, 2004635 (2021).

16. S. Lemma et al., Energy metabolism in osteoclast formation and activity. Int J Biochem Cell Biol 79, 168–180 (2016).

17. B. Li et al., Both aerobic glycolysis and mitochondrial respiration are required for osteoclast differentiation. FASEB J 34, 11058–11067 (2020).

18. H. N. Kim et al., Estrogens decrease osteoclast number by attenuating mitochondria oxidative phosphorylation and ATP production in early osteoclast precursors. Sci Rep 10, 11933 (2020).

19. F. Bost, L. Kaminski, The metabolic modulator PGC-1alpha in cancer. Am J Cancer Res 9, 198–211 (2019).

20. K. A. Ishii et al., Coordination of PGC-1beta and iron uptake in mitochondrial biogenesis and osteoclast activation. Nat Med 15, 259–266 (2009).

21. W. Wei et al., PGC1beta mediates PPARgamma activation of osteoclastogenesis and rosiglitazone-induced bone loss. Cell Metab 11, 503–516 (2010).

22. Y. Zhang et al., PGC1beta Organizes the Osteoclast Cytoskeleton by Mitochondrial Biogenesis and Activation. J Bone Miner Res 33, 1114–1125 (2018).

23. J. Heward et al., KDM5 inhibition offers a novel therapeutic strategy for the treatment of KMT2D mutant lymphomas. Blood 138, 370–381 (2021).

24. L. Wu et al., KDM5 histone demethylases repress immune response via suppression of STING. PLoS Biol 16, e2006134 (2018).

25. D. Lu et al., LRP1 Suppresses Bone Resorption in Mice by Inhibiting the RANKL-Stimulated NF-kappaB and p38 Pathways During Osteoclastogenesis. J Bone Miner Res 33, 1773–1784 (2018).

26. J. Li et al., Desumoylase SENP6 maintains osteochondroprogenitor homeostasis by suppressing the p53 pathway. Nat Commun 9, 143 (2018).

27. H. Guak et al., PGC-1beta maintains mitochondrial metabolism and restrains inflammatory gene expression. Sci Rep 12, 16028 (2022).

28. S. A. Mookerjee, A. A. Gerencser, D. G. Nicholls, M. D. Brand, Quantifying intracellular rates of glycolytic and oxidative ATP production and consumption using extracellular flux measurements. J Biol Chem 292, 7189–7207 (2017).

29. E. H. Ma et al., Metabolic Profiling Using Stable Isotope Tracing Reveals Distinct Patterns of Glucose Utilization by Physiologically Activated CD8(+) T Cells. Immunity 51, 856–870 e855 (2019).

30. B. Cordeiro et al., MicroRNA-9 Fine-Tunes Dendritic Cell Function by Suppressing Negative Regulators in a Cell-Type-Specific Manner. Cell Rep 31, 107585 (2020).

31. G. M. Boukhaled et al., The Transcriptional Repressor Polycomb Group Factor 6, PCGF6, Negatively Regulates Dendritic Cell Activation and Promotes Quiescence. Cell Rep 16, 1829–1837 (2016).

