## Supplemental Data for "The histone demethylase KDM5C controls female bone mass by promoting energy metabolism in osteoclasts"

Science
AAAS

### Supplementary Materials for

#### The histone demethylase KDM5C controls female bone mass by promoting energy metabolism in osteoclasts

Huadie Liu[2]†, Lukai Zhai[1]†, Ye Liu[2], Di Lu[2], Alexandra VanderArk[1], Tao Yang[2]*, and Connie M. Krawczyk[1]*

**This PDF file includes:**

Materials and Methods
Figs. S1 to S5

#### Materials and Methods

##### Mice

The *KDM5C*$^{fl/fl}$ mouse with C57BL/6 background was a kind gift from Yang Shi's lab. In brief, exons 11 and 12 of *Kdm5c* were flanked by *flox* sequences to enable excision by Cre recombinase (*10*). *Kdm5c*$^{fl/fl}$ mice were crossed to *Vav*-iCre (strain 008610) and *LysM*-Cre (strain 004781) mice, respectively, purchased from Jackson Laboratory to generate KDM5C knockout mice. All mice were maintained following guidelines of the Institutional Animal Care and Use Committee (IACUC) at Van Andel Institute. Mice euthanization was conducted according to American Veterinary Medical Association (AVMA) Guidelines for the Euthanasia of Animals.

##### Bone histomorphometry

Bone histomorphometry was performed as described before (*25*). Briefly, paraffin-embedded femurs from two-month-old mice were sent to Pathology and Biorepository Core at Van Andel Research Institute for sectioning. TRAP staining on paraffin sections were performed using an Acid Phosphatase, Leukocyte kit (Sigma-Aldrich, 387A) by following the standard protocol. Osteoclast number/bone surface (N.Oc/BS), osteoclast surface/bone surface (Oc.S/BS), and osteoclast surface/osteoclast number (Oc.S/N.Oc) quantified using BioQuant osteo software (Nashville, TN, USA).

##### Micro-CT

Mice femurs were scanned following the JBMR recommended micro-CT guidelines as previously described (*26*). Reconstruction of cross-section slices was performed using NRecon software (SkyScan) while bone parameters were calculated through CTAN and CTVOL software. Bone parameters calculated are bone volume/total volume (BV/TV), trabecular thickness (Tb.Th), trabecular separation (Tb.Sp), trabecualr number (Tb.N), cortical bone area (Ct.Ar), cortical thickness (Ct.Th), the Ct.Ar/Tt.Ar ratio, and total cross-sectional area (Tt.Ar).

##### Bone marrow monocytes (BMM) isolation and culture

Bone marrow cells were collected from femurs and tibias and plated in complete BMM media (a-MEM plus 10% FBS, 1% pen/strep, 1% L-glutamine, and 30 ng/ml M-CSF). The next day, nonadherent cells were transferred to new dishes for BMM culture with fresh complete BMM media and expanded for 48 hours.

##### Human peripheral blood mononuclear cells (PBMC) extraction and culture

Human blood collection was approved by the Internal Review Board (IRB) of Calvin University (reference number: 20-023). All participating donors have understood and signed IRB-approved consent forms. PBMC were obtained by centrifugation of blood through a Ficoll-Hypaque density gradient at 300×g for 60 min. Adherent cells from PBMC were cultured in BMM media for 1-3 days before use.

##### *Ex vivo* osteoclastogenesis assay

BMMs or cultured human monocytes ($3\times10^4$ cells/ well or $6\times10^4$ cells/ well respectively) were seeded into 96-well plates and cultured overnight with complete BMM media. Differentiation was induced by osteoclastogenic medium (50 ng/ml RANKL in complete BMM media) and induction media was changed every other day. Cells were subjected to staining when osteoclast fusion appeared (3-5 days for BMMs,7-9 days for human monocytes). TRAP staining was conducted to visualize osteoclasts using an Acid Phosphatase Leukocyte (TRAP) Kit (Sigma-Aldrich) by following the standard protocol. Cells positive for TRAP staining and contain > 3 nuclei were counted as osteoclasts. Osteoclast numbers and area were quantified using ImageJ.

**Total RNA isolation and qRT-PCR**

Total RNA from BMMs were extracted using TRIzol reagent (Invitrogen) by following the standard protocol. Genomic DNAs were removed by using the DNA-free™ DNA Removal kit (ThermoFisher, AM1906) according to the manufacturer's instruction. 500 ng RNA were subjected to the synthesis of first-strand cDNA using SuperScript™ VILO™ cDNA synthesis Kit (Invitrogen, 11754050). Quantitative PCR (qPCR) was performed on a StepOne PCR instrument using SRYBR Green QPCR Master Mix (Invitrogen, 4472908). Primers used for qRT-PCR in this study are: *Kdm5c*: 5'-GAGCAGTCTGTACTGTGCCA-3' (forward), 5'-ATTCCCACATACAGCCACGG-3' (reverse); *Nfatc1*: 5'-TCATCGGCGGGAAGAAGATG-3' (forward), 5'-GTCCCGGTCAGTCTTTGCTT-3' (reverse); *Fos*: 5'-TACTACCATTCCCCAGCCGA-3' (forward), 5'-GCTGTCACCGTGGGGATAAA-3' (reverse); *Oscar*: 5'-CGTGCTGACTTCACACCAAC-3' (forward), 5'-GGTCACGTTGATCCCAGGAG-3' (reverse); *Dcstamp:* GCTGTATCGGCTCATCTCC*T*-3' (forward), 5'-ATGGACGACTCCTTGGGTTC-3' (reverse), *; Ocstamp:* 5'-AGCCACGGAACACCTCTTTG-3' (forward), 5'-TGGAACAACTGCCTTGCAGA-3' (reverse)*;* *Ctsk*: 5'-GAAGGGAAGCAAGCACTGGA -3' (forward), 5'-CCATGTTGGTAATGCCGCAG-3' (reverse); *Acp5*: 5'-CTGGTATGTGCTGGCTGGAA-3' (forward), 5'-CGCAAACGGTAGTAAGGGCT-3' (reverse); *Atp6v0d2*: 5'-GGGCCTGGTTCGAGGATG-3' (forward), 5'-GAAGTTGCCATAGTCCGTGGT-3' (reverse); *Uqcrc2*: 5'-GCAACTGCTAGAGCCATGAAG-3' (forward), 5'-TTAACCTTCGGGGCAACTTTGA-3' (reverse); *Apt1b3*: 5'-TGCTGGAACCAGGAACCTAAA-3' (forward), 5'-CTAGGCTCGTGCTGTGACTT-3' (reverse); *Cox6b1*: 5'-CGCTACTCCGGGACAATCTT-3' (forward), 5'-TCTGGTTCTGGTTGGGGAAG-3' (reverse); *Sdhb*: 5'-CGTTCTCGGCAGAGTCGG-3' (forward), 5'-GGTCCCATCGGTAAATGGCA-3' (reverse); *Sdha*: 5'-TATGGTGCAGAAGCTCGGAAG-3' (forward), 5'-ACTCATCGACCCGCACTTTG-3' (reverse); *Ppargc1b*: 5'-CAGTACAGCCCCGATGACTC (forward), 5'-TTCGTAAGCGCAGCCAAGA-3' (reverse); *Ppia*: 5'-AGCATACAGGTCCTGGCATC-3' (forward), 5'- TTCACCTTCCCAAAGACCAC -3' (reverse). *Ppia* gene was used as the internal control for normalization. The mRNA levels of genes of interest were normalized to the average levels of *Ppia,* relative expression was calculated as the ratio to control (defined as 1).

**RNA sequencing and data processing**

BMM from fCtrl and f*Kdm5c*$^{\Delta Vav}$ (n=3) mice were seeded into 12-well plates followed by RANKL treatment for 0 hours (untreated), 16 hours or 32 hours. Total RNA was extracted using the same method as described in qRT-PCR and sequenced by the Genomics Core at Van Andel Research Institute. Raw reads of RNA-seq data were mapped to *Mus musculus* (mm10). Subjunc v1.6.4 and featureCounts v1.6.4 were used to estimate the read counts on transcripts/gene exons. EdgeR v3.32.1 were used to identify differentially expressed genes. Genes with FDR ≤ 0.05 (Benjamini-Hochberg adjusted p-values) were annotated as a differentially expressed gene between fCtrl and f*Kdm5c*$^{\Delta Vav}$ BMM. R package enrichR v3.1 was used to identify gene sets (Gene Ontology Biology Process 2021) enriched in the differentially expressed genes. Terms with p-value <0.05 were considered significant. The top 5 terms ranked by combined score of up- and down-regulated genes are listed in Fig 4A. Combined score is computed using the logarithm of Fisher exact test p-value multiplex by z-score.

**Western blots**

Cultured BMM were lysed in CHAPS buffer (Thermo Scientific) containing protein inhibitors. 10-40 μg protein per sample were loaded to 10% SDS PAGE gels followed by transfer to PVDF membranes. The membranes were then blocked with 5% non-fat milk in 1× TBST (Tris-buffered saline plus 0.05% Tween® 20) for two hours at room temperature and incubated with corresponding primary antibodies. Primary antibodies: anti-KDM5C (A301-034A, Bethyl Laboratories) or anti-β-actin (4967, Cell Signaling Technology), OxPhos Rodent WB Antibody Cocktail (45-8099, Thermo Scientific). Blots were developed using SuperSignalt™ West Dura Extended Duration Substrate (Thermo Scientific) and imaged using ChemiDoc™ MP Imaging System (BIO-RAD).

**Extracellular flux assay**

Seahorse assay and ATP calculations were performed as described (*27–29*). Oxygen consumption rate (OCR) and extracellular acidification rate (ECAR) of cultured BMM were measured using the Seahorse XF96 Extracellular Flux Analyzer. In brief, twenty thousand BMM were seeded to XF96 plates and treated with RANKL to induce osteoclastogenesis. Three days after RANKL treatment, cellular bioenergetics were assessed using Seahorse XF Cell Mito Stress Test Kit with sequential addition of 1.5 μM Oligomycin (Oligo: inhibiting ATP synthesis), 3 μM carbonyl cyanide 4-(trifluoromethoxy) phenylhydrazone (FCCP: uncoupling), 0.5 μM rotenone/antimycin A (Rot/AA: inhibiting respiratory chain complex l and lll, respectively), and 10 mM monensin (Mon: stimulating Na+ pumps on plasma membrane). Data were normalized to number of cells.

**BMM retroviral transfection**

BMM were transduced with control (pMSCV-ires-Thy1.1; pMIT) or PGC1-b expressing (pMiT-*Pgc1-β*) retrovirus as previously described (*30*, *31*). Briefly, 293T cells were transfected with vectors and Lipofectamine 2000 to generate retroviral supernatants. BMM were cultured in 6-well plates with complete BMM media until 70% confluency was reached. Cells were then transfected with pMiT or pMiT-*Pgc1-β* retrovirus containing supernatants by centrifuging under 2500 rpm for 60 min at 30 °C, prior to being cultured in fresh complete BMM media. Two days after transfection, cells were digested with 0.5 mM EDTA (Ethylenediaminetetraacetic acid), stained with PE-Thy-1.1 antibody (12-0900-81, Thermo Fisher SCIENTIFIC) and sorted using anti-PE microbeads (Miltenyi Biotec). Sorted BMM were seeded in 96-well plates for osteoclastogenesis, in 12-well plates for RNA extraction and western blots, or in XF96 plates for Seahorse assays.

**KDM5 inhibitor treatment**

The pan-KDM5 inhibitor, KDM5A-IN-1, (KDM5i) was purchased from MedChemExpress (Cat. No.: HY-100014). The half maximal effective concentration (EC50) was defined as concentration at which the osteoclast numbers and area are halved compared to the control group. Cultured BMM from fCtrl mice or human monocytes were thereafter treated with DMSO or KDM5i for osteoclastogenesis assay, Seahorse assay or RNA extraction.

**Statistical analysis**

All data were analyzed using GraphPad Prism software (version 9). An unpaired or paired Student's *t*-test was performed for experiments with two groups for statistical significance calculation. A one-way or two-way ANOVA analysis of variance was performed to determine statistical significance between multiple groups. Actual P values are shown on each graph.

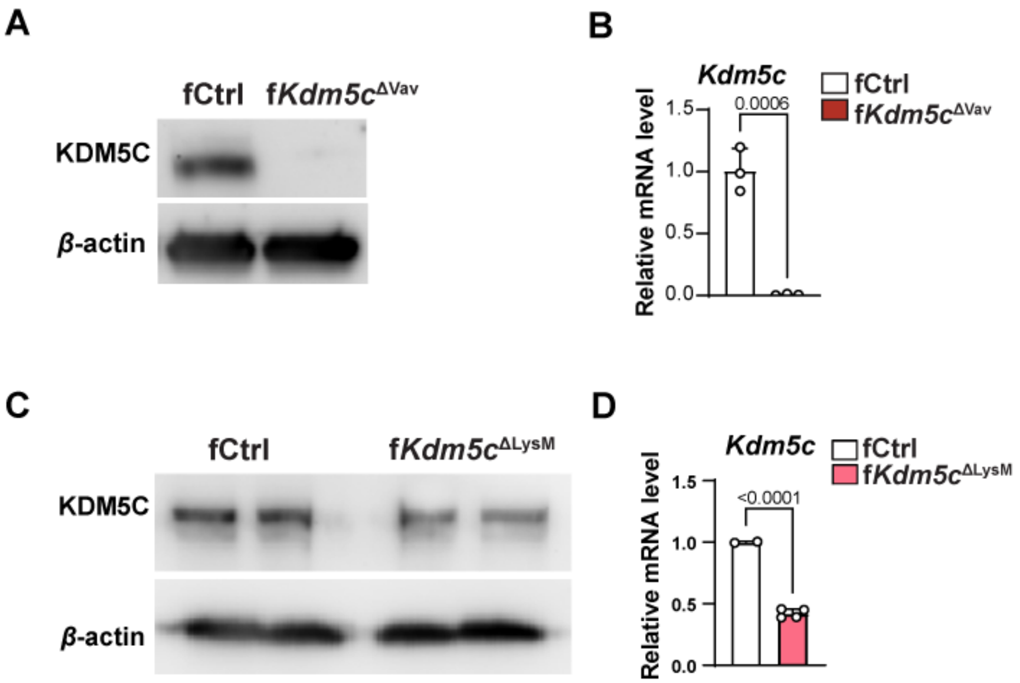


**Fig. S1.**

**Reduced deletion efficiency by *LysM-Cre* compared to *Vav-Cre* in BMM.** (**A**-**B**) Protein levels (A) and mRNA levels (B) of KDM5C in f*Kdm5c*$^{\Delta Vav}$ and fCtrl BMM. (**C**-**D**) Protein levels (C) and mRNA levels (D) of KDM5C in f*Kdm5c*$^{\Delta LysM}$ and fCtrl BMM. All data comparisons are conducted by Student's *t*-test, two-tailed. Data are presented as mean ± s.e.m. .

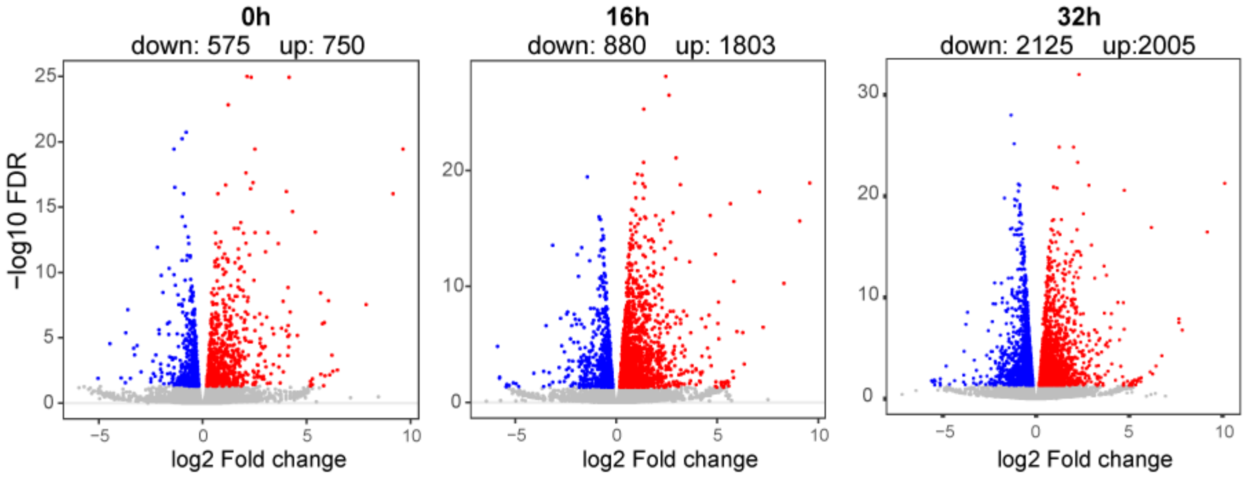


**Fig. S2.**

**Transcriptomic changes between f*Kdm5c*$^{\Delta Vav}$ and fCtrl BMM during *ex vivo* osteoclastogenesis assay.** Volcano plot of RNA-seq data showing significant changes in gene expression between f*Kdm5c*$^{\Delta Vav}$ and control BMMs at different osteoclastogenesis stages (0h, 16h, and 32h after RANKL stimulation).

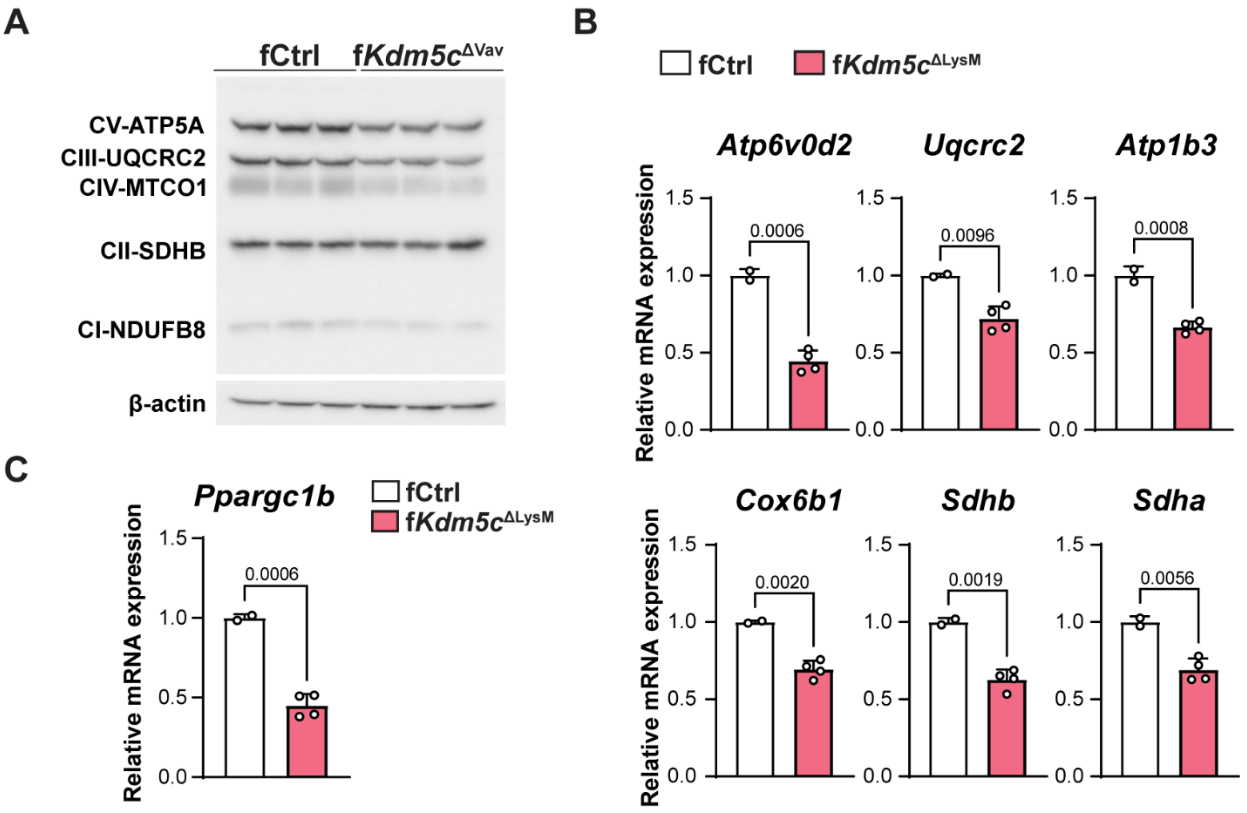


**Fig. S3.**

***Ppargc1b* and mitochondrial respiration genes are repressed in KDM5C-deficient BMM.**
(**A**) Lower exposure of Fig.4E of mitochondrial OXPHS complex proteins in f*Kdm5c*$^{\Delta Vav}$ and fCtrl BMMs (**B**) mRNA levels of mitochondrial respiration genes in f*Kdm5c*$^{\Delta LysM}$ and fCtrl BMM. (**C**) Level of *Ppargc1b* mRNA in f*Kdm5c*$^{\Delta LysM}$ and fCtrl BMM. All data comparisons are conducted by Student's *t*-test, two-tailed. Data are presented as mean ± s.e.m.

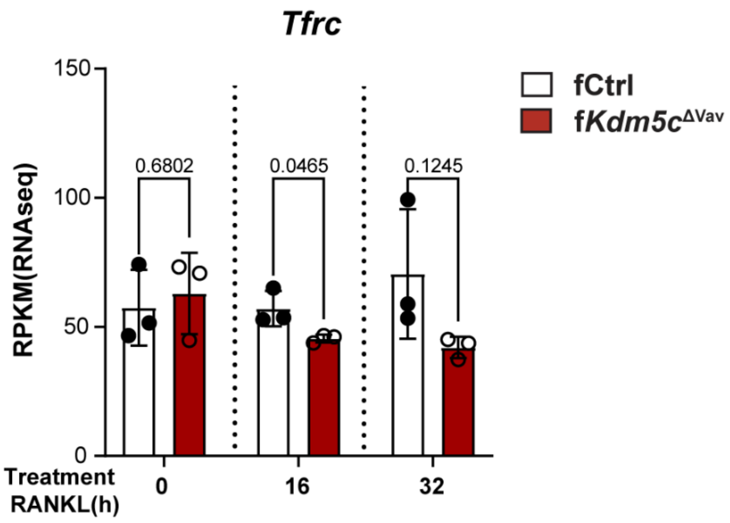


**Fig. S4.**

**KDM5C-deficeint BMMs have decreased transferrin receptor gene expression during osteoclastogenesis.** Level of Transferrin receptor protein 1 gene (*Tfrc1*) transcripts in fCtrl and f*Kdm5c*$^{\Delta Vav}$ BMM during osteoclastogenesis (Student's *t*-test, two-tailed). Data are presented as mean ± s.e.m.

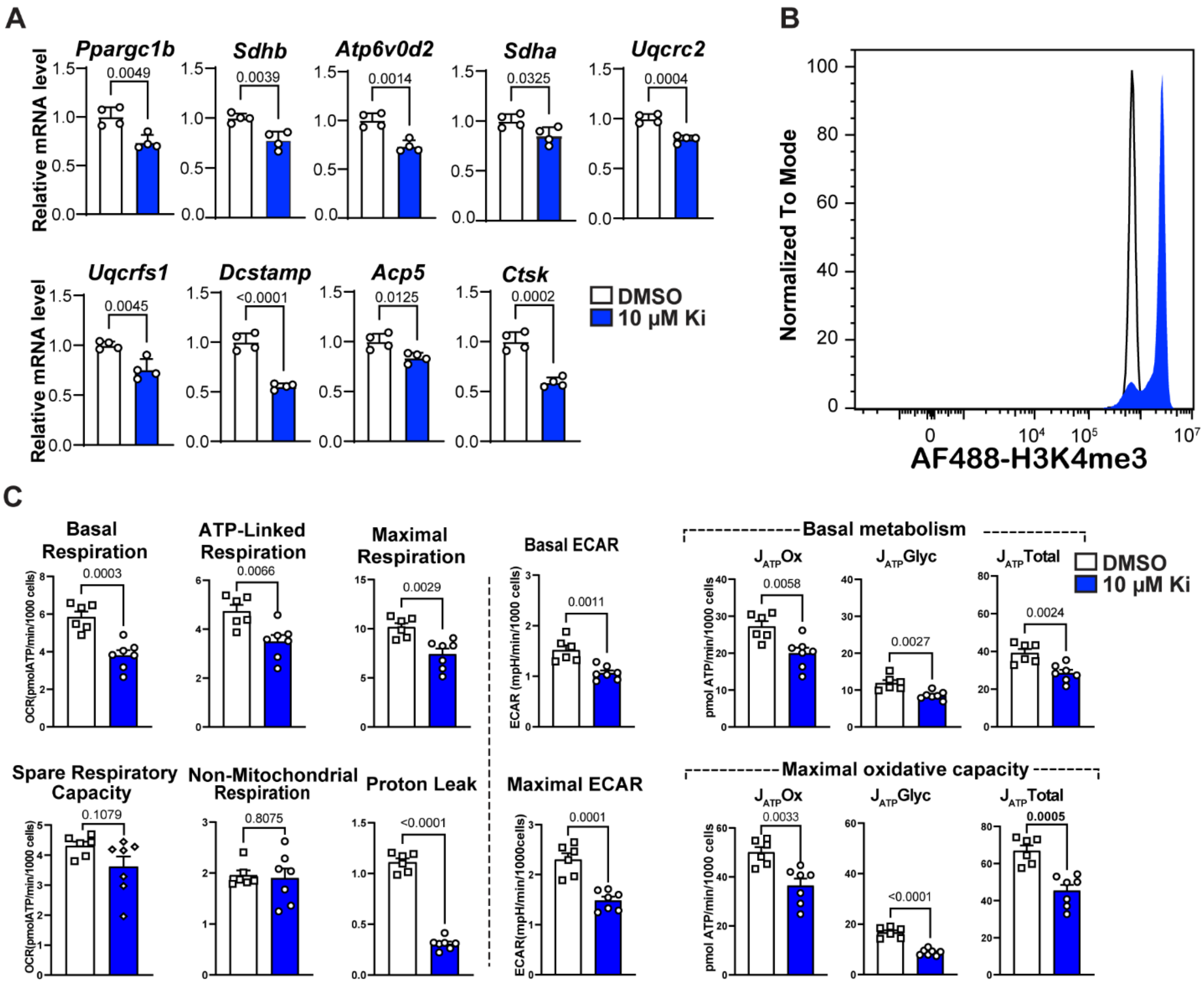


**Fig. S5.**

**Pharmacological inhibition of KDM5 increases H3K4me3 and impairs mitochondrial metabolism and osteoclastogenesis of mouse and human monocytes.** (**A**) Expression of *Ppargc1b,* osteoclast and mitochondrial respiration genes in DMSO and Ki treated mouse BMMs. mRNA was detected by qRT-PCR 48h after RANKL treatment. (**B**) H3K4me3 levels in control and Ki treated CD14+ monocytes cultured from human peripheral blood. H3K4me3 levels were detected by flow cytometry 24h after inhibitor treatment. (**C**) Detailed parameters of OCR, ECAR and Oxidative/glycolytic/total ATP production in control and Ki-treated human PBMC-monocyte 5 days after osteogenic induction. Comparisons in (A and C) are conducted by Student's *t*-test, two-tailed. Data are presented as mean ± s.e.m.
